# *Plasmodium falciparum* Research in Africa from 2000 to 2024: A Systematic Review and Bibliometric Visualisation

**DOI:** 10.1101/2025.10.12.681826

**Authors:** Dogara Elisha Tumba, Dauda Wadzani Palnam, Peter Abraham, Israel Ogwuche Ogra, Samson Usman, Morumda Daji, Dasoem Naanswan Joseph, Mela Ilu Luka, Jonathan Utos Emohchonne, Ndukwe K. Johnson, Grace Peter Wabba, Seun Cecilia Joshua, Mercy Nathaniel, Elkanah Glen, Zainab Kasim Mohammed, Umezuruike Linus Opara

## Abstract

*Plasmodium falciparum* is the major cause of malaria in Africa, responsible for high morbidity and mortality across the continent. This study presents a systematic literature review and bibliometric analysis of *P. falciparum* research conducted in Africa between 2000 and 2024 as shown in Figure 1. Using the PRISMA framework, 10,903 peer-reviewed articles were retrieved from the Scopus database. Bibliometric analysis was performed using Bibliometrix in R and VOSviewer to assess publication trends, authorship networks, keyword evolution, and thematic concentration. Results reveal contributions from 18,345 authors across 4,903 journals, with an average of 31.83 citations per article. Research output has grown steadily over the two-decade period, with significant input from African scholars and international collaborators. The most active research themes include epidemiology, antimalarial drug resistance, vaccine development, vector biology, and socio-economic factors in malaria control. Despite this progress, the review highlights persistent gaps in genomic surveillance, localised insecticide resistance monitoring, and integration of social determinants into malaria intervention strategies. Regional disparities in research output remain, with some high-burden areas underrepresented. Collaboration among African institutions is limited compared to international partnerships. These observations indicate the urgent need for targeted funding, strengthened intra-African collaboration, and policies that contextualise malaria research within local health systems. Addressing these gaps is essential for speeding the continent’s malaria elimination agenda.

Graphical representation and summary of trends in *Plasmodium falciparum* in Africa.

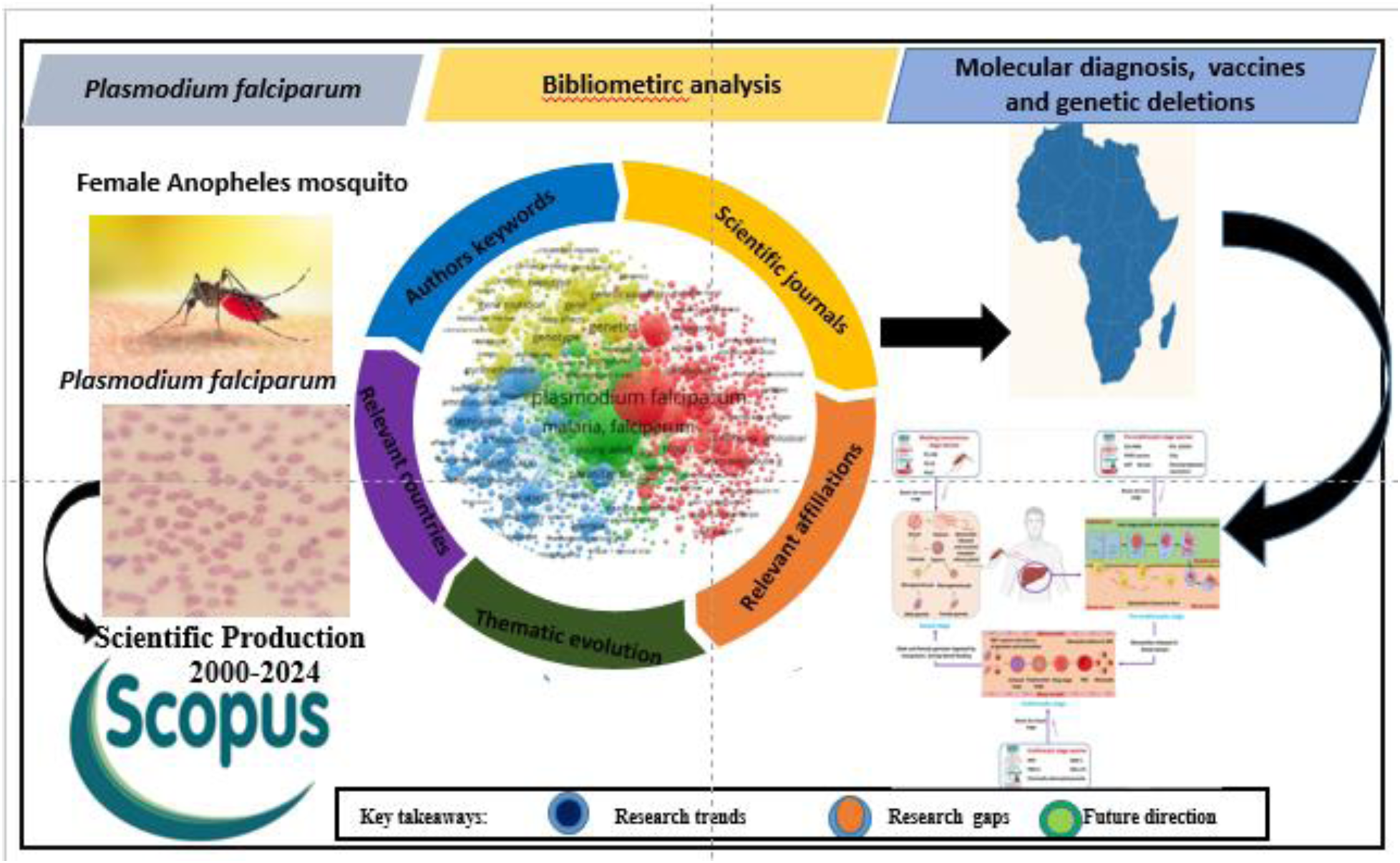

## 1.0 INTRODUCTION

Malaria is a life-threatening disease caused by several species of the Plasmodium parasite, including *Plasmodium falciparum*, *Plasmodium vivax*, *Plasmodium malariae*, *Plasmodium ovale*, and *Plasmodium knowlesi*. Among these, *Plasmodium falciparum* and *Plasmodium vivax* are known to contribute most significantly to the global burden of malaria-related morbidity and mortality (WHO, 2023). This alarming impact is underscored by data from the World Health Organization’s 2023 malaria key facts, which report an estimated 249 million cases and 608,000 deaths in 85 endemic countries. Notably, the WHO African Region bears the heaviest toll, accounting for 94% of global cases and 95% of deaths, with children under the age of five being the most vulnerable demographic (WHO, 2023). The transmission of malaria occurs through the bite of infected female mosquitoes of the *Anopheles* genus, which inject Plasmodium parasites directly into the human bloodstream.

Of particular concern is *Plasmodium falciparum*, one of the six protozoan species infecting humans, and widely recognized as the most virulent and deadly. It is responsible for approximately 80% of human malaria cases globally and is especially prevalent across tropical Africa, where environmental conditions favor its persistence and spread (Cowman *et al.,* 2016).

Given its dominant role in malaria pathology, *Plasmodium falciparum* has become the focal point of many systematic reviews aiming to synthesize existing literature on malaria in Africa. These reviews offer a broad lens through which to view research developments, consolidate findings, and inform policy-making and future scientific inquiry. By leveraging basic research, collaborative networks, systematic reviews, and bibliometric analyses, malaria research continues to evolve, driving innovations that enhance malaria control and eradication efforts across Africa. Specifically, they provide a bibliometric and systematic overview of research outputs, highlight key thematic areas, and identify gaps in knowledge surrounding diagnostic strategies and interventions (Oyegoke *et al.,* 2022).

Beyond cataloging published studies, these reviews also direct attention toward underexplored dimensions such as asymptomatic malaria and the socio-economic factors driving incidence rates (Worrall *et al.,* 2002; Asmelash *et al*., 2025). Bibliometric analysis plays a critical role here by uncovering emerging topics, methodological shifts, and geographic patterns in malaria scholarship. This in turn facilitates the mapping of collaborative networks and the recognition of leading institutions contributing to the field (Phoobane *et al.,* 2022; Chutiyami, 2024).

Finally, citation analysis and related metrics serve as valuable tools to evaluate the impact of malaria research. They reveal which studies have shaped the scientific discourse, gauge knowledge dissemination, and help prioritize high-impact investigations. Collectively, these analytic approaches ensure that future research is both strategic and contextually relevant, thereby strengthening the fight against malaria across Africa (Chutiyami, 2024).

This review aims to systematically analyze *Plasmodium falciparum* research conducted in Africa between 2000 and 2024, using bibliometric visualization to identify prevailing trends, research hotspots, and gaps in knowledge. It seeks to evaluate the evolution of scientific output, highlight influential studies and collaborations, and provide evidence-based insights to guide future research priorities and malaria control strategies across the continent.

To achieve this objective, the following research questions were proposed as a guide for this study: Research Question 1: How has scientific production evolved in research on *Plasmodium falciparum* in Africa?

Research Question 2: What are the trending and major themes on *Plasmodium falciparum* research in Africa?

Research Question 3: What are the research gaps and future directions on *Plasmodium falciparum research in Africa?*

## 2.0 METHODOLOGY

### 2.1 Research Design

This study adopted a bibliometric analysis of published research studies from 2000 to 2024. As an emerging field in scientific research, bibliometric analysis enables researchers to systematically explore, collect, and analyze large volumes of scientific data to identify key landmarks and emerging research trends in a specific field (Donthu *et al*., 2021). Oliveira et al. (2019) added that bibliometric analysis offers insight into the performance of authors, articles, journals, institutions, and countries. This study adheres to key performance indicators and science mapping outlined by Donthu *et al*. (2021), presenting a systematic bibliometric analysis guideline.

For the retrieval of datasets to be used for bibliometric analysis and systematic review, various databases and search engines were used. These include: Scopus, ScienceDirect, Google Scholar, PubMed, Proquest and PsycINFO. The number of documents retrieved from the various “<u>databases</u>” are presented in Table 1,

**Table 1:** Number of Publications related to *Plasmodium falciparum* in Africa retrieved from Different Database.

| SN | SEARCH ENGINE OR DATABASE | NUMBER OF PAPER RETRIEVED |
| --- | --- | --- |
| 1 | Scopus | 11,014 |
| 2 | Google scholar | 17,500 |
| 3 | PubMed | 4,162 |
| 4 | Science direct | 9,029 |
| 5 | Proquest | 20,058 |
| 6 | PsycINFO | 67,472 |
|  | Retrival date | 17<sup>th</sup>, November 2024. |

### 2.2 Data Collection

The dataset was retrieved on August 26, 2024 at about 2:02PM from the Scopus database, known for its extensive coverage of academic publications. The search strategy utilized specific search strings within the “Title-Abstract-Keywords” fields. These strings contained a combination of Boolean operators (OR and AND), truncation signs (*), and quotation marks (“. . .”) to ensure comprehensive and relevant retrieval. The search strings used for this study are (“*Plasmodium falciparum*” OR “*P. falciparum*”) AND (“Africa”)

### 2.3 Inclusion and Exclusion Criteria

The search yielded a broad spectrum of articles, including those in English and published before the data retrieval date. This step was essential to ensure the analysis was based on the latest and most relevant literature. A manual review of the identified records was then conducted to exclude documents not specifically related to *Plasmodium falciparum* or falciparum malaria. The final step involved exporting the records meeting our inclusion criteria in .csv format for bibliometric analysis (Figure 1). This process was crucial for building a robust dataset for our study. From 10,903 retrieved documents, only 4,903 publications were considered eligible for further analysis

**Figure 1:**
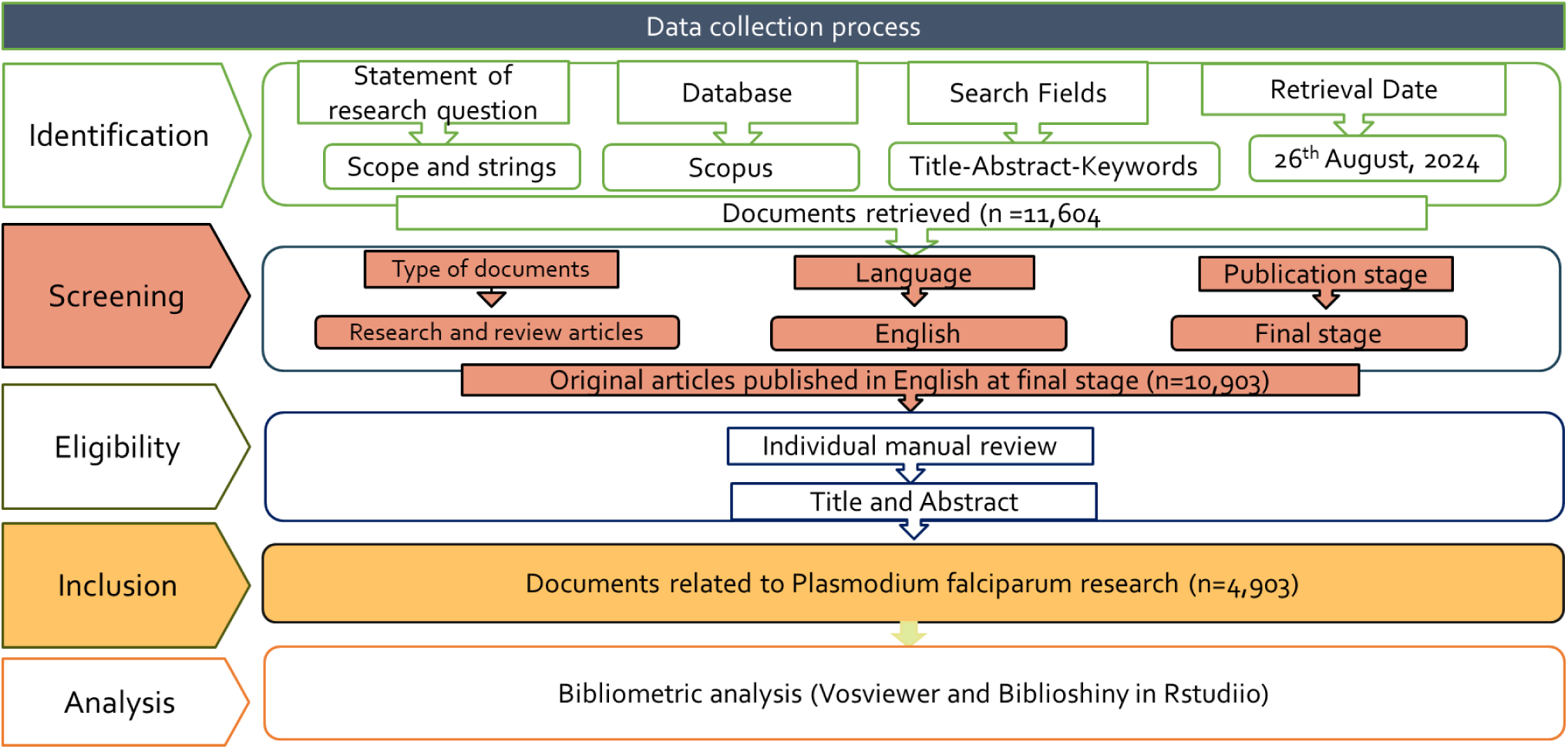
Flow chart of Methodology using PRISMA method.

### 2.4 Analysis and Visualization

We utilized data visualization and analysis software like VOSviewer and Biblioshiny to conduct various analyses on the included literature, including co-authorship network analysis, co-occurrence analysis, and thematic word analysis. Bibliometrix facilitated the illustration of scientific trends and productivity within the documents, identifying the most prolific authors and significant articles published on the subject. This package encompasses powerful and comprehensive capabilities for bibliometric analysis, comprising analyses of authors, institutions, countries, and regions, as well as journal clustering and temporal trends. In contrast, VOSviewer offers an intuitive approach to analyzing knowledge within research fields, supporting various visualization views to facilitate a deeper understanding of research trends and theme development.

## 3.0 RESULTS

### 3.1 Overview of Research Statistics on *Plasmodium falciparum* Research in Africa

A total of 10903 publication outputs on *Plasmodium falciparum* were identified in the Scopus database. (Figure 1) The global scientific production on *Plasmodium falciparum* research consists of 4903 documents (Figure 2). The production comprises papers published in English from 2000 to 2024 in the Scopus database (10,903). They are strictly related to *Plasmodium falciparum* research. Furthermore, these 4903 research articles have been authored by 18345 contributors across 665 sources with an average growth citation rate of 31.83. The annual growth rate shows that *Plasmodium falciparum* research appears to be increasing over time with 1.98%. The international collaboration of 87.78% highlights a significant degree of international networking in *Plasmodium falciparum* research in Africa. Out of the 10903 publications, 31 articles are with single authors. The distribution of document types on global research on Plasmodium falciparum research in Africa The 4903 documents used for the analysis are distributed as articles (96%) and reviews(4%) in Figure. 3

**Figure 2:**
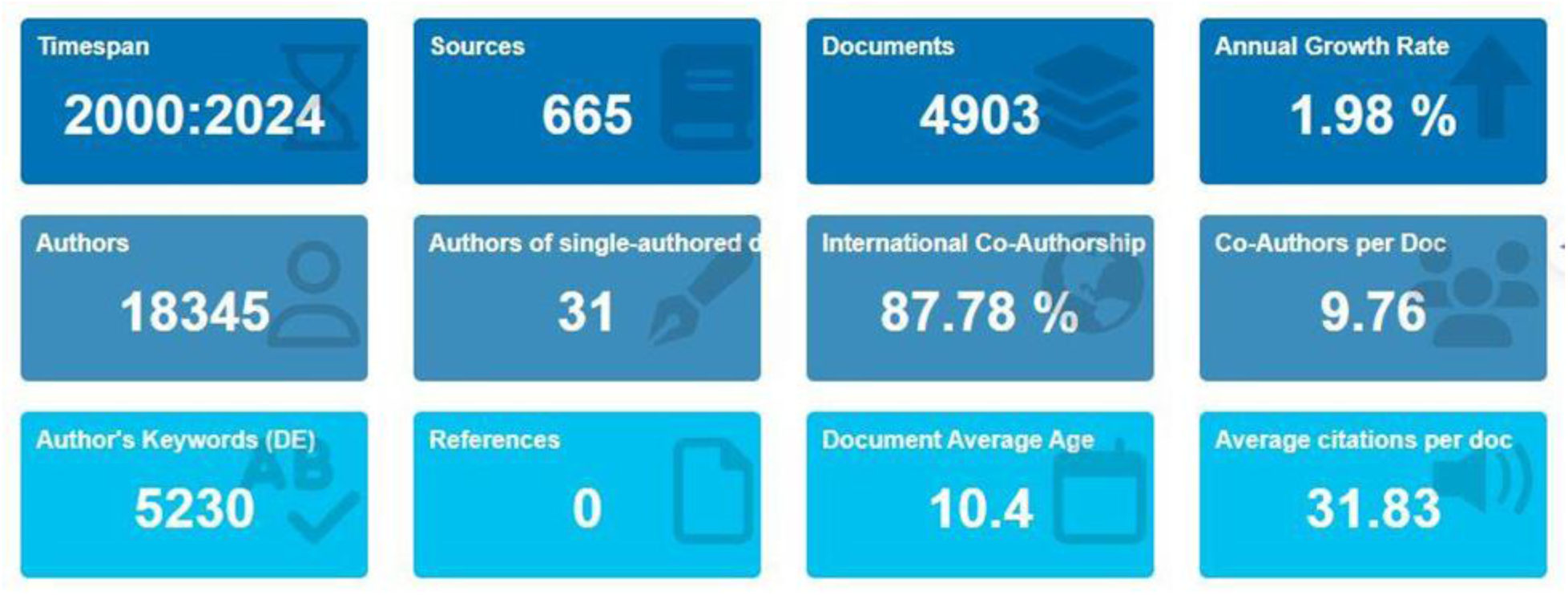
An Overview of Research performance statistics on *Plasmodium falciparum* research in Africa based on Scopus.

**Figure 3:**
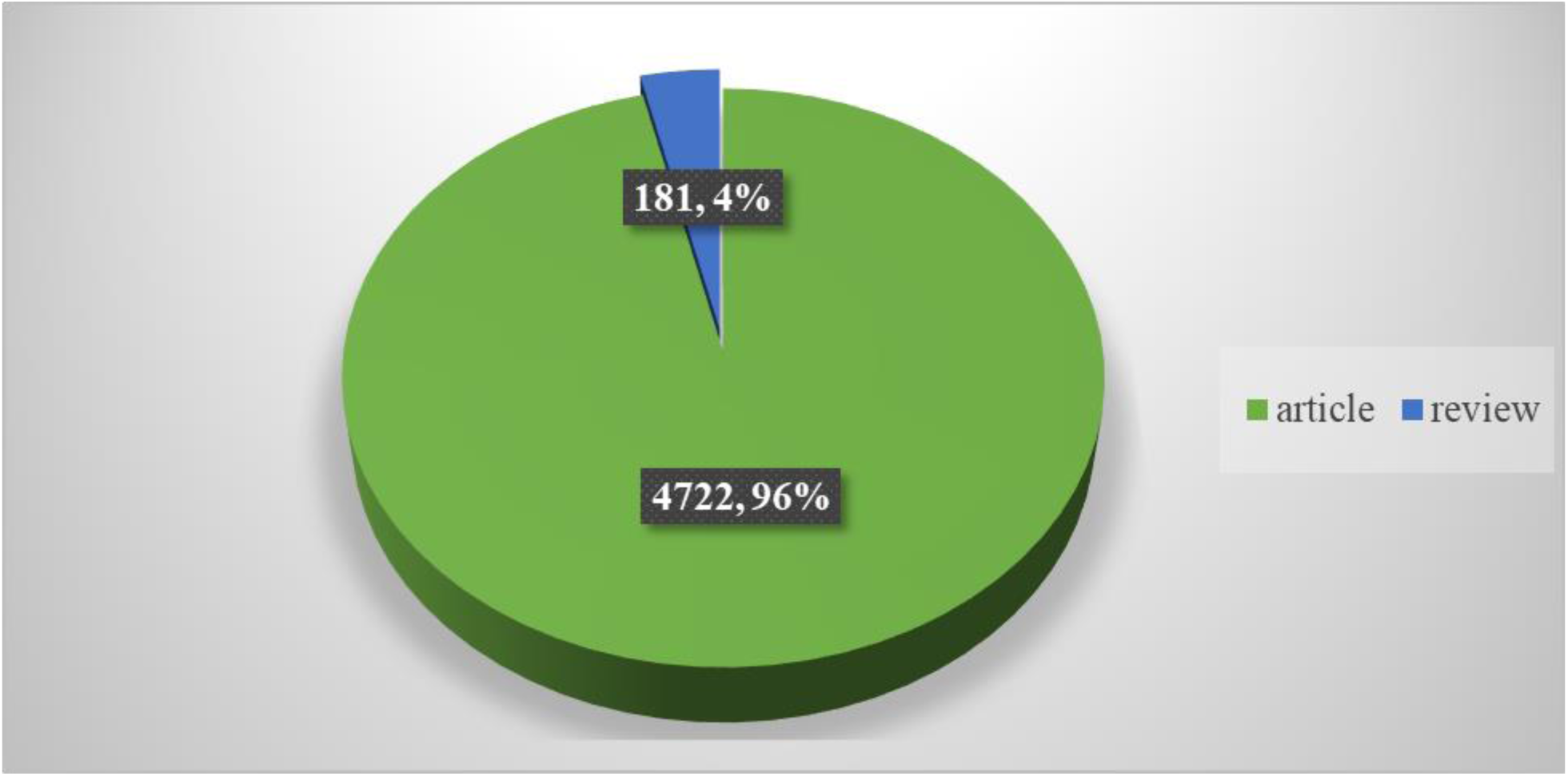
Distribution of Document Types on *Plasmodium falciparum* Research in Africa.

### 3.2 Publication Trends on *Plasmodium falciparum* research in Africa

#### 3.2.1 Research Publication Trends

The annual production rate of articles on *Plasmodium falciparum* in Africa is shown in Figure 4. There has been a gradual increase in publication from 2001 to 2021. The highest increase in publication on *Plasmodium falciparum* research in Africa was observed between 2021 and decline in 2022 to 2024. Figure 5 showed that the highest mean citation per articles was recorded in the years 2014 to 2016.

**Figure 4:**
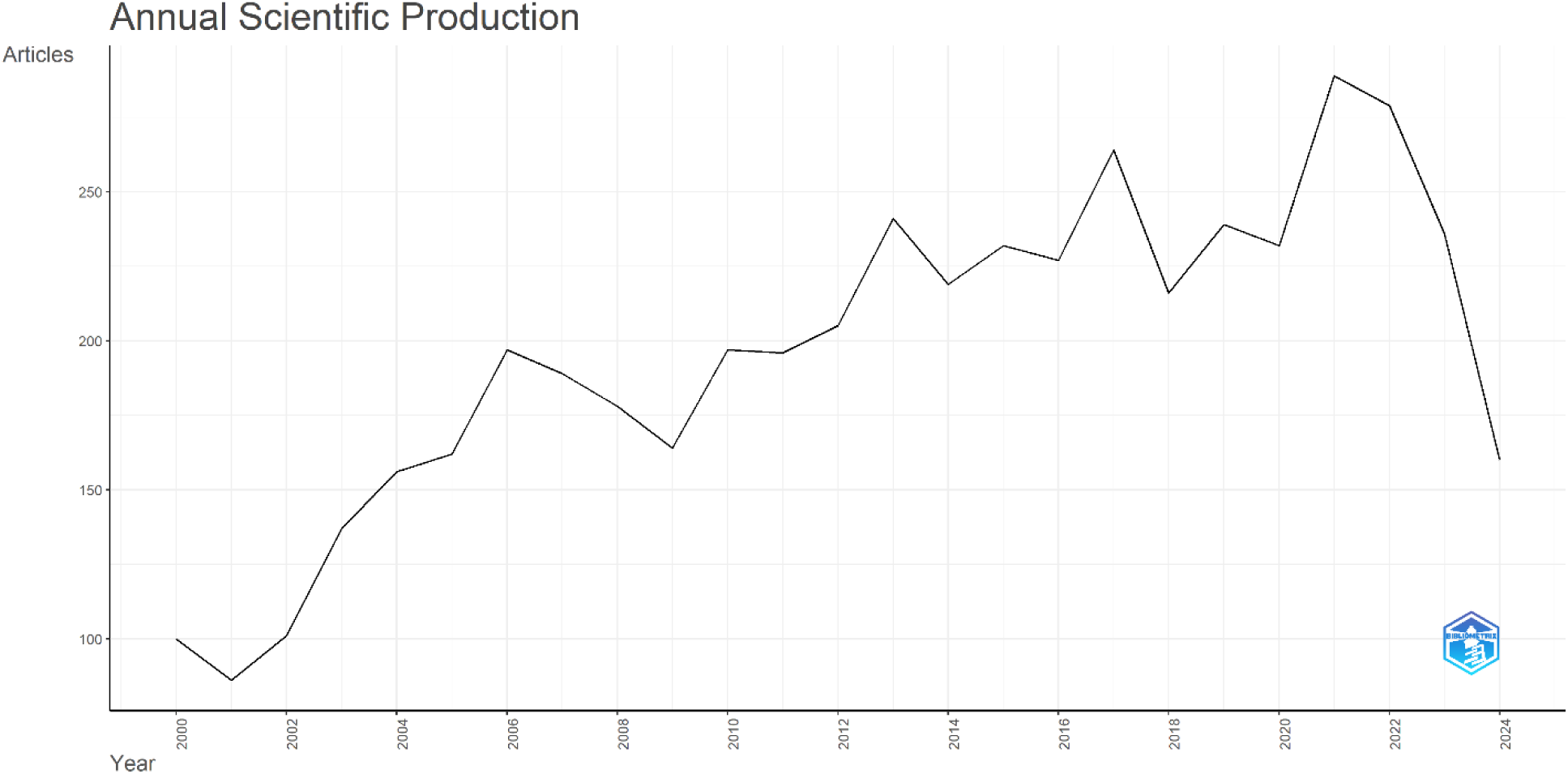
Annual Scientific Production of articles on *Plasmodium falciparum* research in Africa.

**Figure 5:**
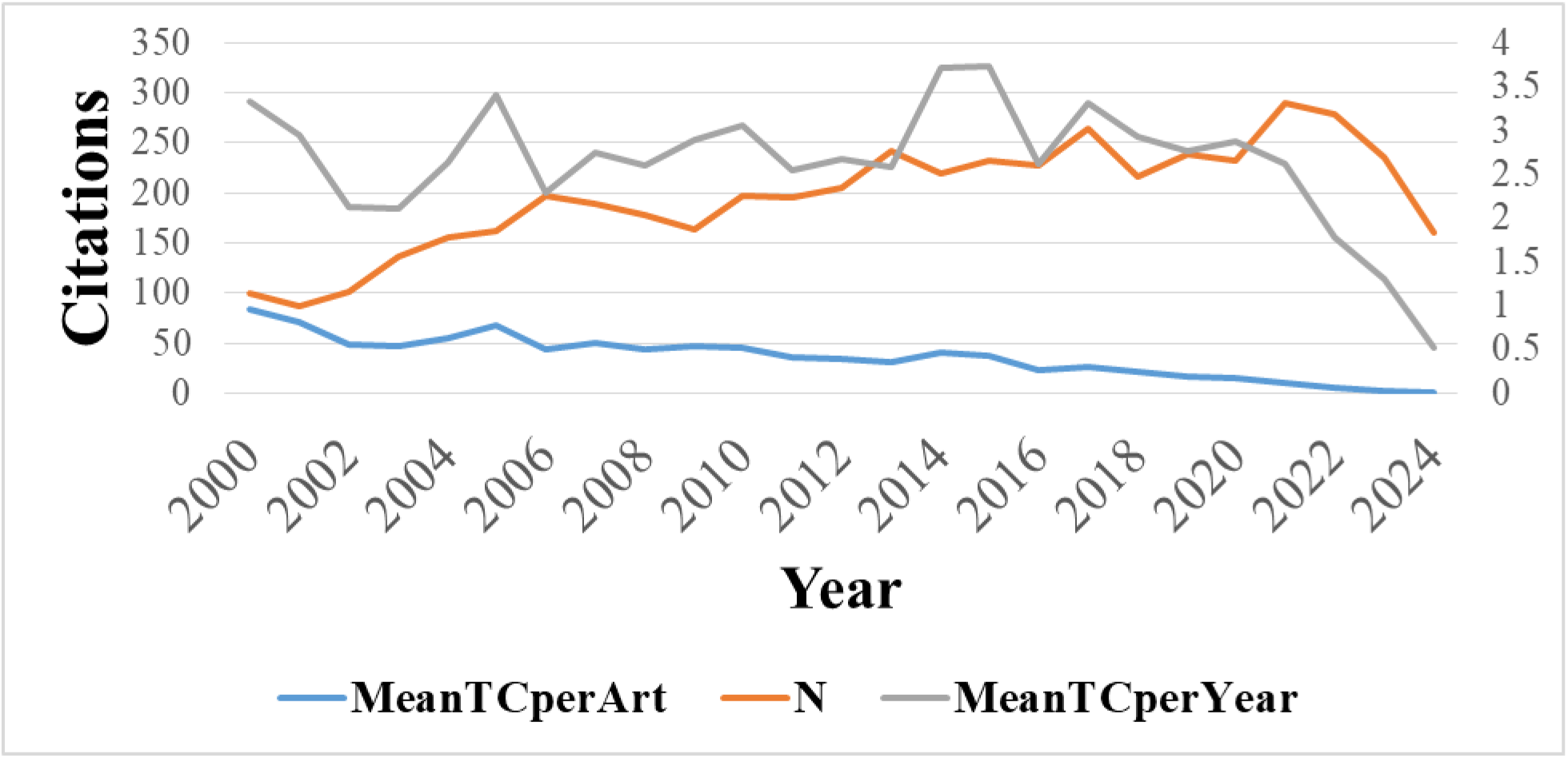
Total Citation per Year of Documents on *Plasmodium falciparum* Research in Africa.

### 3.3 Global Collaboration Network on Sickle Cell Disease Research in Africa

The global collaboration network in *Plasmodium falciparum* research in Africa is characterized by partnerships from different countries, thus facilitated by shared research interest in *Plasmodium falciparum* research in Africa (Figure 6). African countries like Nigeria, Kenya, South Africa and Ghana are actively contributing to malaria research with significant documents counts with emphasis on the need to address the malaria burden in Africa, where *Plasmodium falciparum* is prevalent. The inter-country collaboration involves developed nations such as the USA, UK and France.

**Figure 6:**
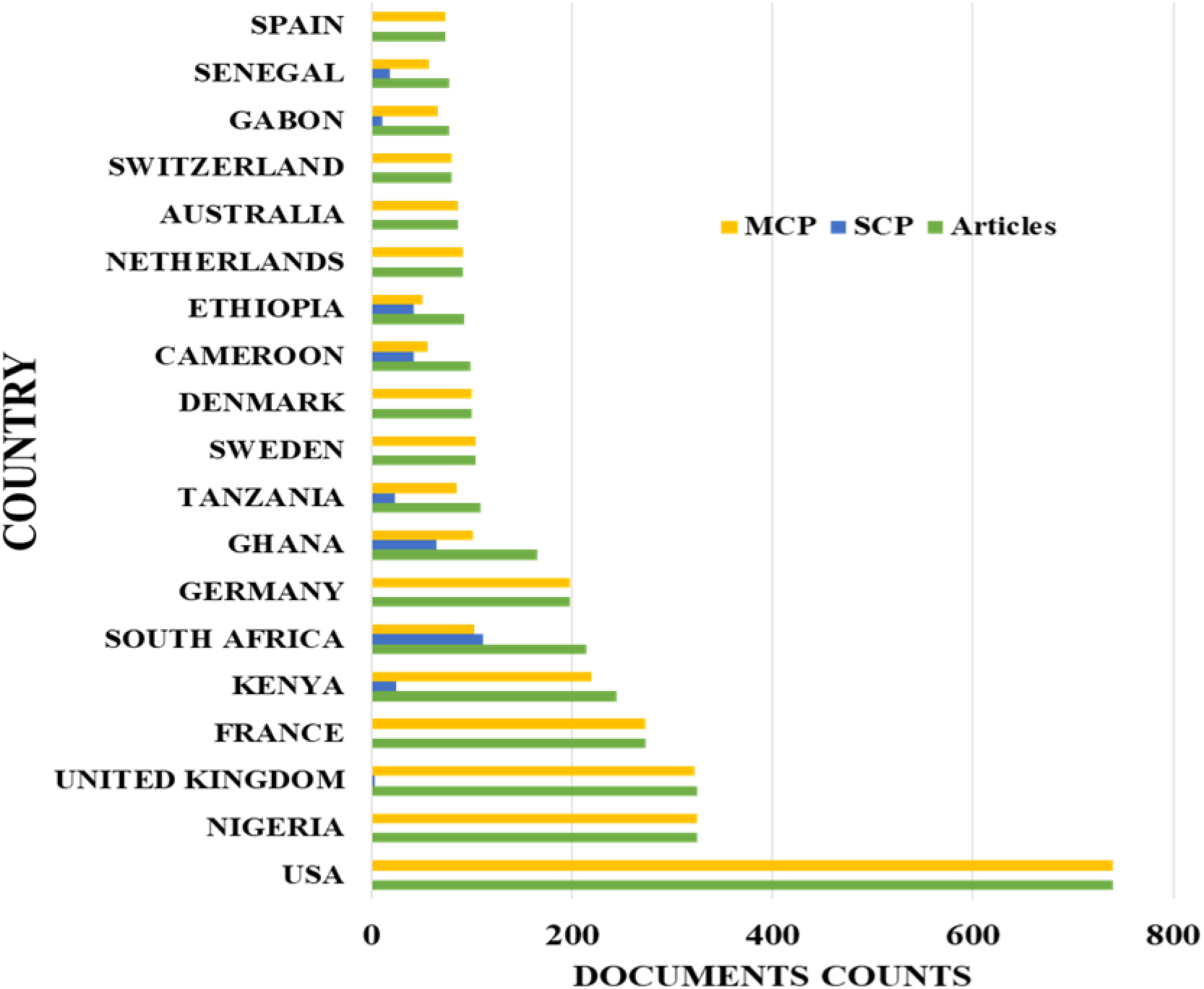
Corresponding Author’s Countries SCP and MCP Represent Intra-country and Inter-country. Collaborations or Country Publications and Multiple-country Publications, Respectively. SCP: Single Country Publication (Intra-country collaborations); MCP: Multiple Countries Publication (Inter-country collaborations).

The global research on *Plasmodium falciparum* in Africa is represented by contributions from different countries and the top 20 countries that were actively involved in this field of research are shown in Figure 7. Countries with notable publications are the United States with 9980 publications followed by Kenya (n=6146) and the least was Australia (n=1189). This shows that despite the research being in Africa, there has been a lot of collaboration and funding on *Plasmodium falciparum* research from the United States. Furthermore, based on total citations per country (Figure 8), countries like the USA (n=29318), the United Kingdom (n=17847), and Kenya (n=12395) emerged as the most cited, showing significant impact, influence, and relevance in *Plasmodium falciparum* research in Africa.

**Figure 7:**
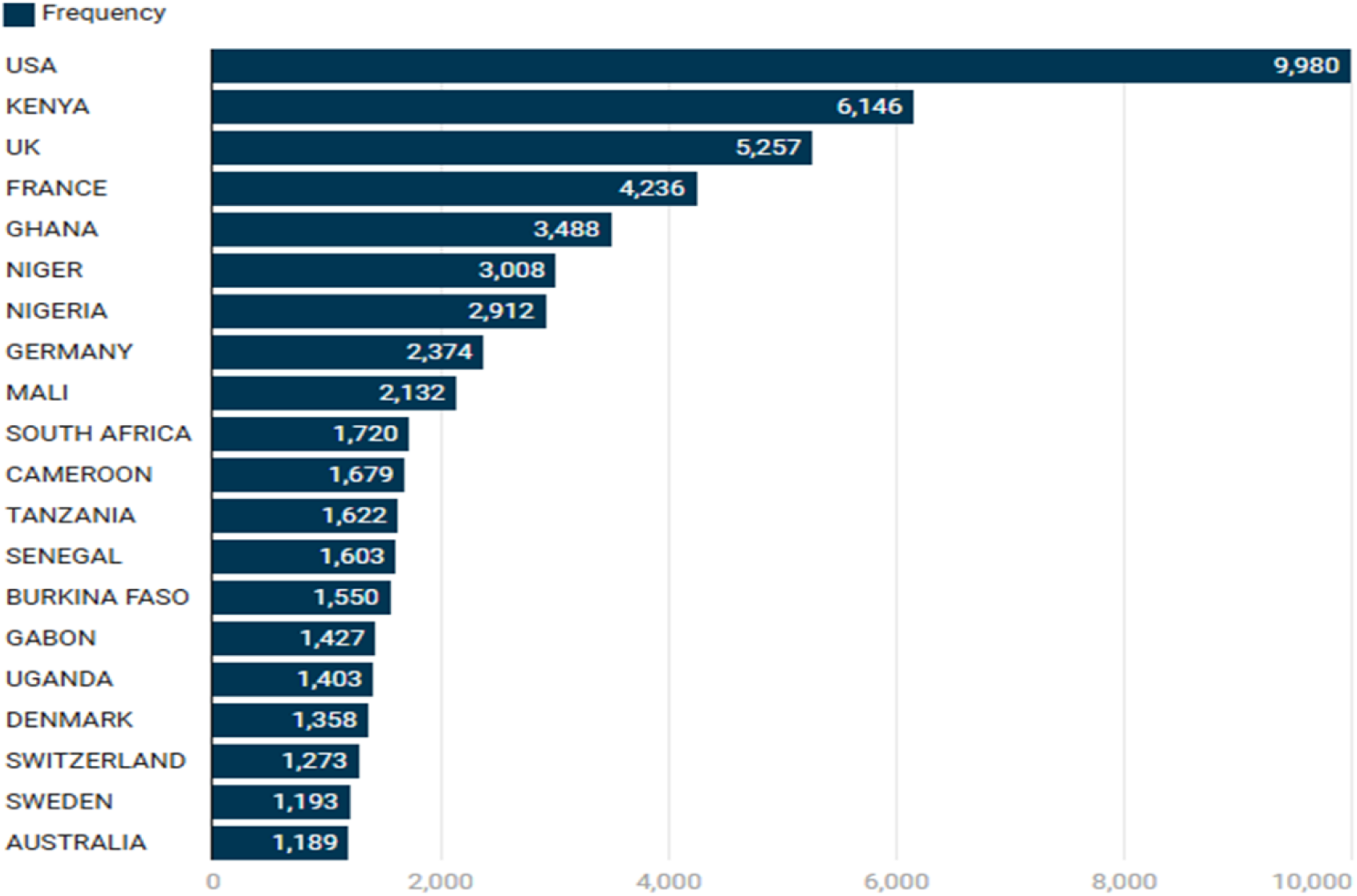
Number of Publications by Top 20 Countries on *Plasmodium falciparum* research in Africa.

**Figure 8:**
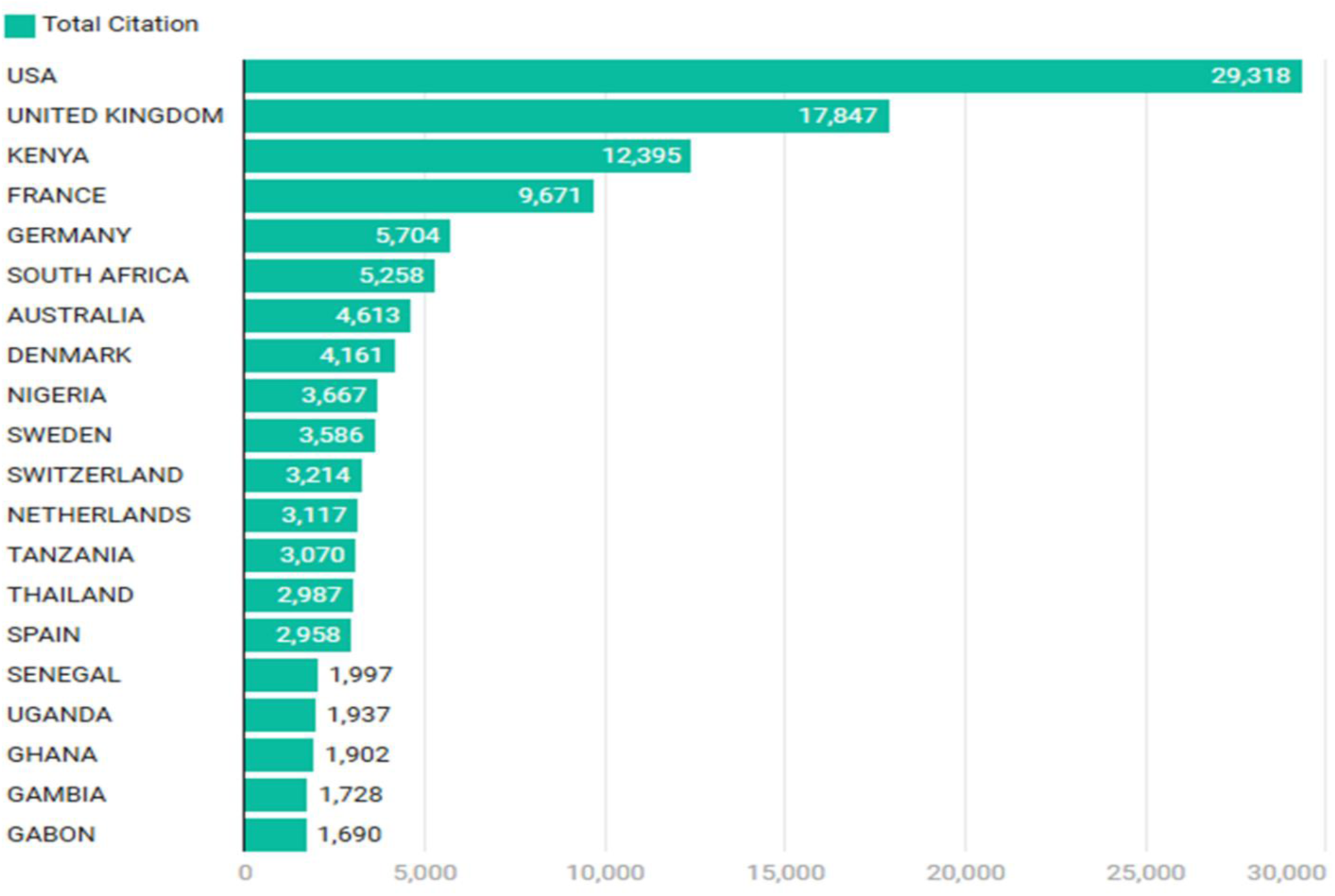
Number of Citations by Top 20 Countries on Global Research on Plasmodium falciparum research in Africa.

### 3.4 Descriptive Statistics of Academic Journals and Most cited Authors of Research on *Plasmodium falciparum* research in Africa

Figure 9 shows the top 10 most productive institutions on *Plasmodium falciparum* research in Africa. The London School of Hygiene is the most productive institution with 911 published articles, followed by the University of Ghana with 873 articles, the University of Oxford with 807 articles, Kenya Medical with 667 articles, and the University of Ibadan with 575 articles. Some institutions in Africa have so many articles on *Plasmodium falciparum because* it is the most prevalent in Africa.

**Figure 9:**
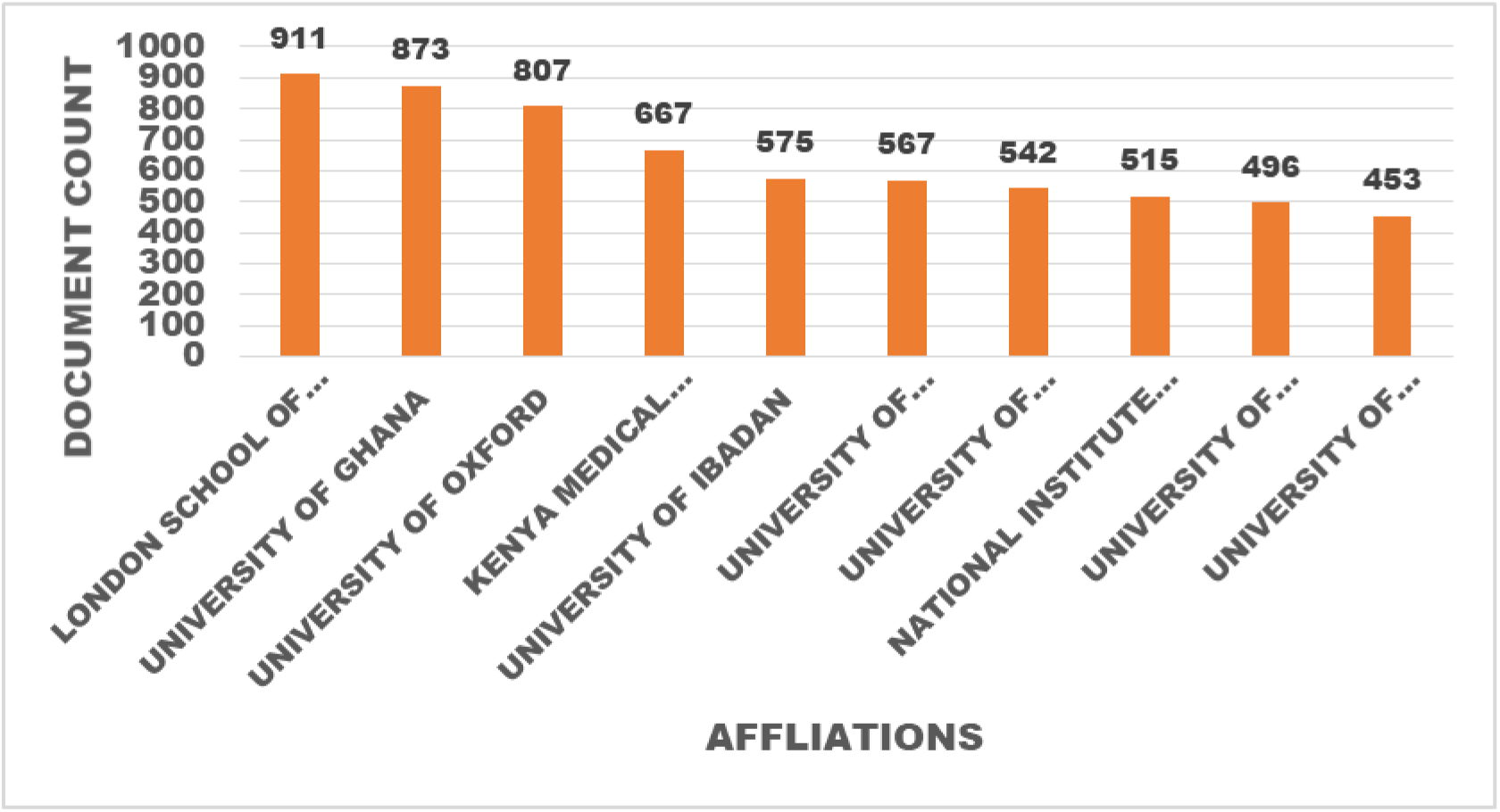
Top 10 Most Relevant Affiliation on *Plasmodium falciparum* research in Africa.

**Figure 10:**
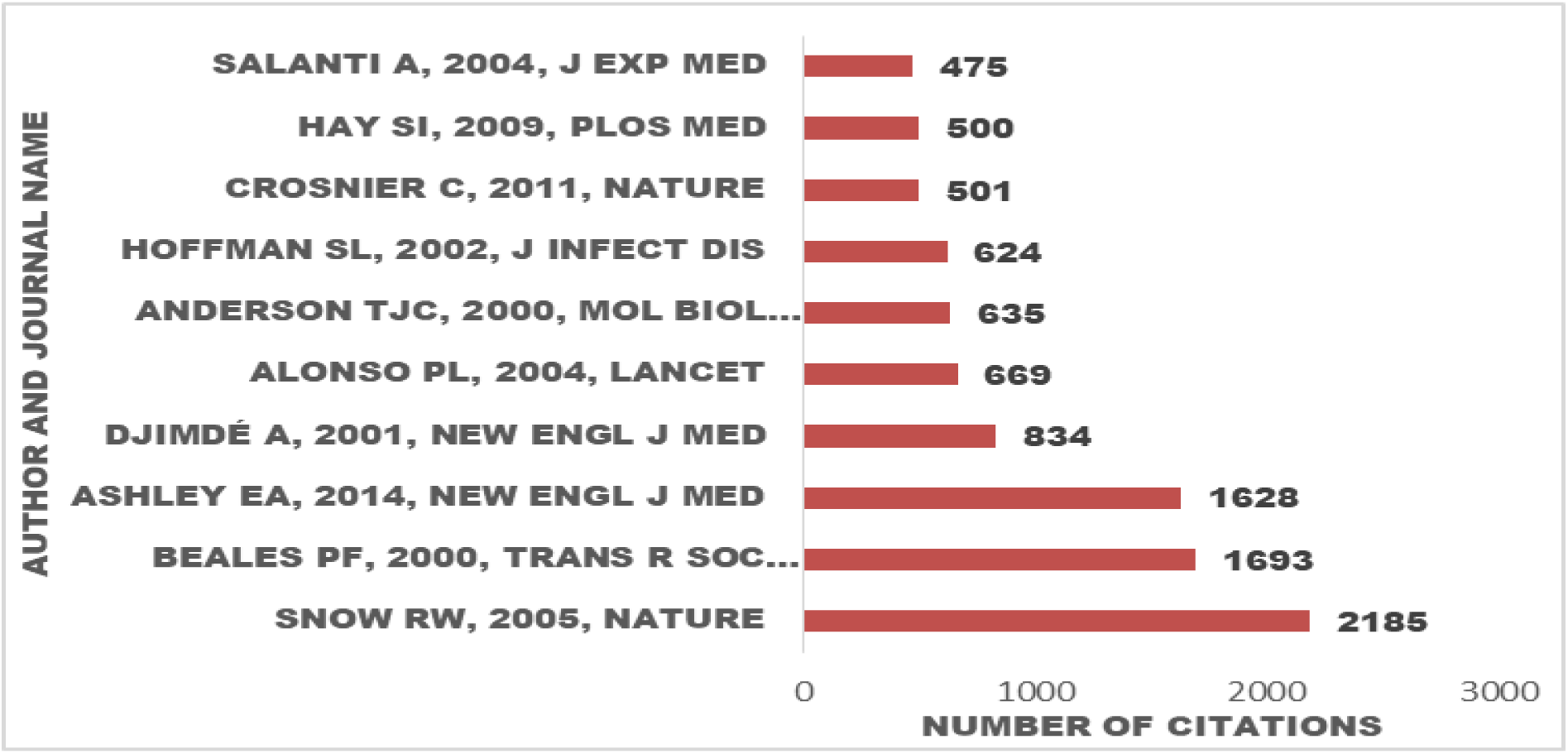
The Top 10 most relevant authors on *Plasmodium falciparum* research in Africa.

**Figure 11:**
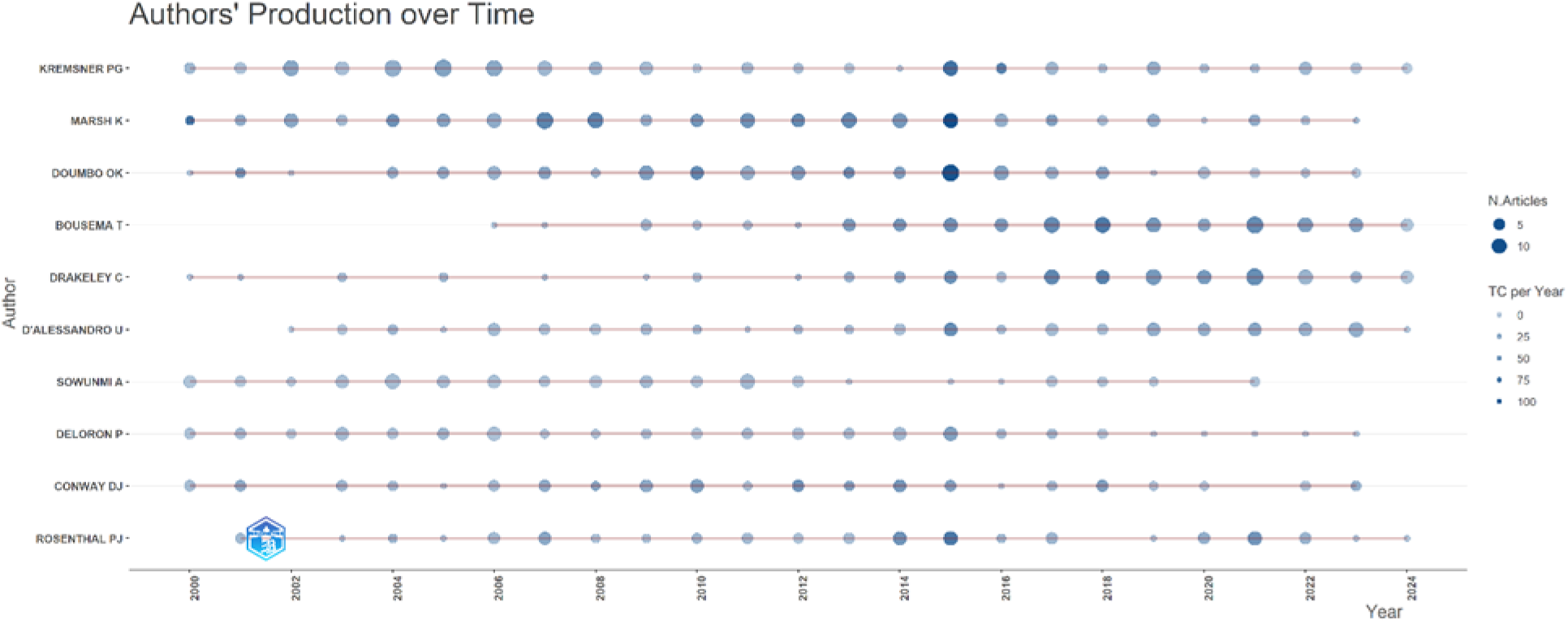
Top authors’ productivity on *Plasmodium falciparum* research in Africa over from 2000 to 2024.

Table 2. shows the top 10 journals in which articles on *Plasmodium falciparum* research in Africa are published. The Journal of Infectious Diseases is the most impactful journal with a h-index of 66 and a total citation of 12984 from 227 publications, followed by Malaria Journal with a h-index of 60 and a total of 19955citations from 880 publications.

**Table 2:** Top 10 Journals in which on *Plasmodium falciparum* are published.

| **S\N**<br>**o** | **Journal** | **h-**<br>**index** | **Number of**<br>**publication** | **Total citations** |
| --- | --- | --- | --- | --- |
| **1** | **JOURNAL OF INFECTIOUS DISEASES** | **66** | **227** | **12984** |
| **2** | **MALARIA JOURNAL** | **60** | **880** | **19955** |
| **3** | **AMERICAN JOURNAL OF TROPICAL**<br>**MEDICINE AND HYGIENE** | **53** | **303** | **9719** |
| **4** | **PLOS ONE** | **49** | **259** | **8118** |
| **5** | **INFECTION AND IMMUNITY** | **44** | **116** | **5628** |
| **6** | **ANTIMICROBIAL AGENTS AND**<br>**CHEMOTHERAPY** | **39** | **142** | **4374** |
| **7** | **TRANSACTIONS OF THE ROYAL SOCIETY**<br>**OF TROPICAL MEDICINE AND HYGIENE** | **36** | **104** | **5153** |
| **8** | **PROCEEDINGS OF THE NATIONAL**<br>**ACADEMY OF SCIENCES OF THE UNITED**<br>**STATES OF AMERICA** | **32** | **34** | **4189** |
| **9** | **TROPICAL MEDICINE AND**<br>**INTERNATIONAL HEALTH** | **30** | **75** | **2473** |
| **10** | **CLINICAL INFECTIOUS DISEASES** | **28** | **55** | **2466** |

**Table 3:**
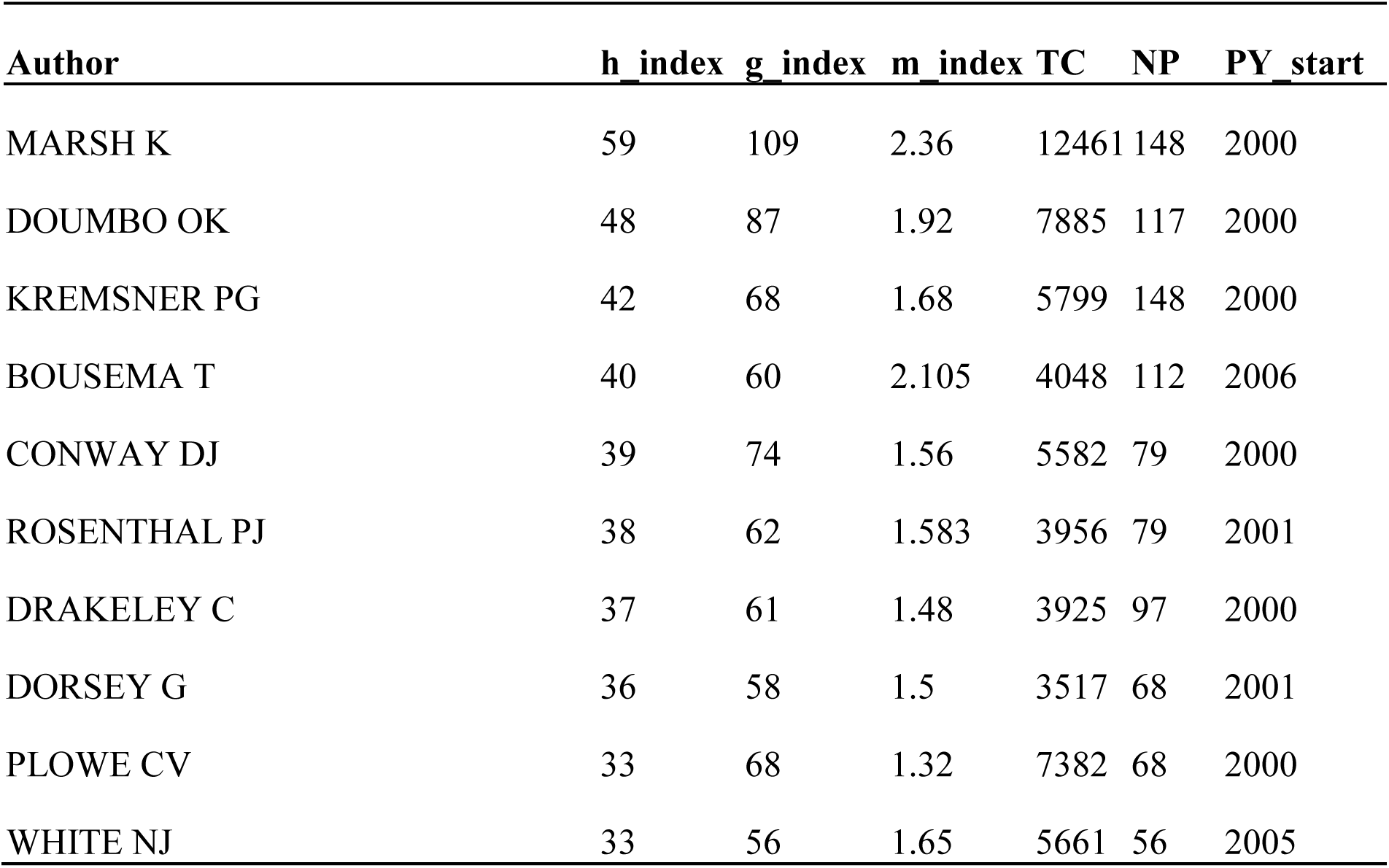
Top 10 Authors in *Plasmodium falciparum* Research in Africa.

### Top 10 Relevant Sources in Global Research on Plasmodium falciparum research in Africa Based on Publications Counts and Impact Factor (h-index)

The most frequent sources include the Malaria Journal, American Journal of Tropical Medicine and Hygiene and PLUS ONE reflecting a significant volume of research on *Plasmodium falciparum* in Africa (Figure 12A). High impact Journals such as the Journal of Infectious Diseases, Infection and Immunity, and Proceedings of the National Academy of Sciences are Prominent. These journals have a strong influence in the field even if they are published in fewer articles than those in Figure 12A.. The dominance of the Malaria Journal and the American Journal of Tropical Medicine and Hygiene suggests that much of the malaria research is published in specialized journals rather than other biological or medical journals. The presence of journals with high-impact factors in Figure 12B highlights the importance of ensuring that malaria research in Africa is published in widely cited and reputable journals. This information is crucial for researchers deciding where to publish their work for maximum visibility and impact.

**Figure 12:**
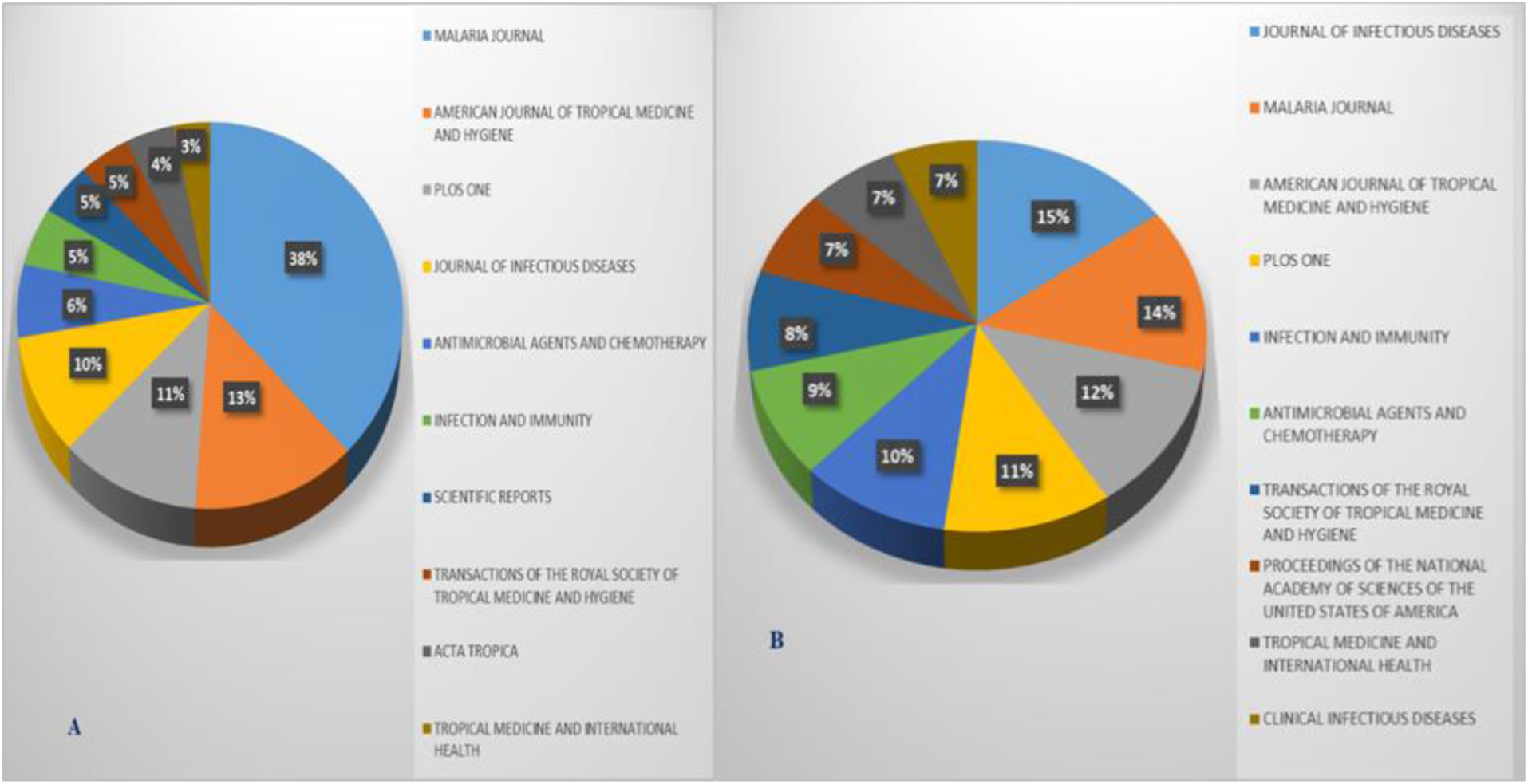
Top 10 Relevant Sources in Global Research on *Plasmodium falciparum* research in Africa. (A) Based on Publications Counts; (B) Based on Impact Factor (h-index)

### 3.5: Thematic Maps of Authors Keywords used in *Plasmodium falciparum* research in Africa

Figure 13 shows the thematic map based on author’s keywords used in *Plasmodium falciparum* research in Africa. It shows the relationship between different keywords used in *Plasmodium falciparum* research in Africa. There are four quadrants; the basic theme, located at the bottom right quadrant, consists of Keywords such as ‘protozoan’ and ‘antigen ‘. These themes have low density but high centrality. The motor theme, positioned at the upper right quadrant, represents the current research focus on *Plasmodium falciparum* in Africa. These themes have high density with low centrality. No keyword was situated in the niche theme (Upper left quadrant). Located at the top left quadrant is the niche theme which represents specialized or less explored areas of research on *Plasmodium falciparum* in Africa. Keywords such as ‘artemisinin’, ‘clinical trials and ‘chloroquine drug resistance’ are found in this region.

**Figure 13:**
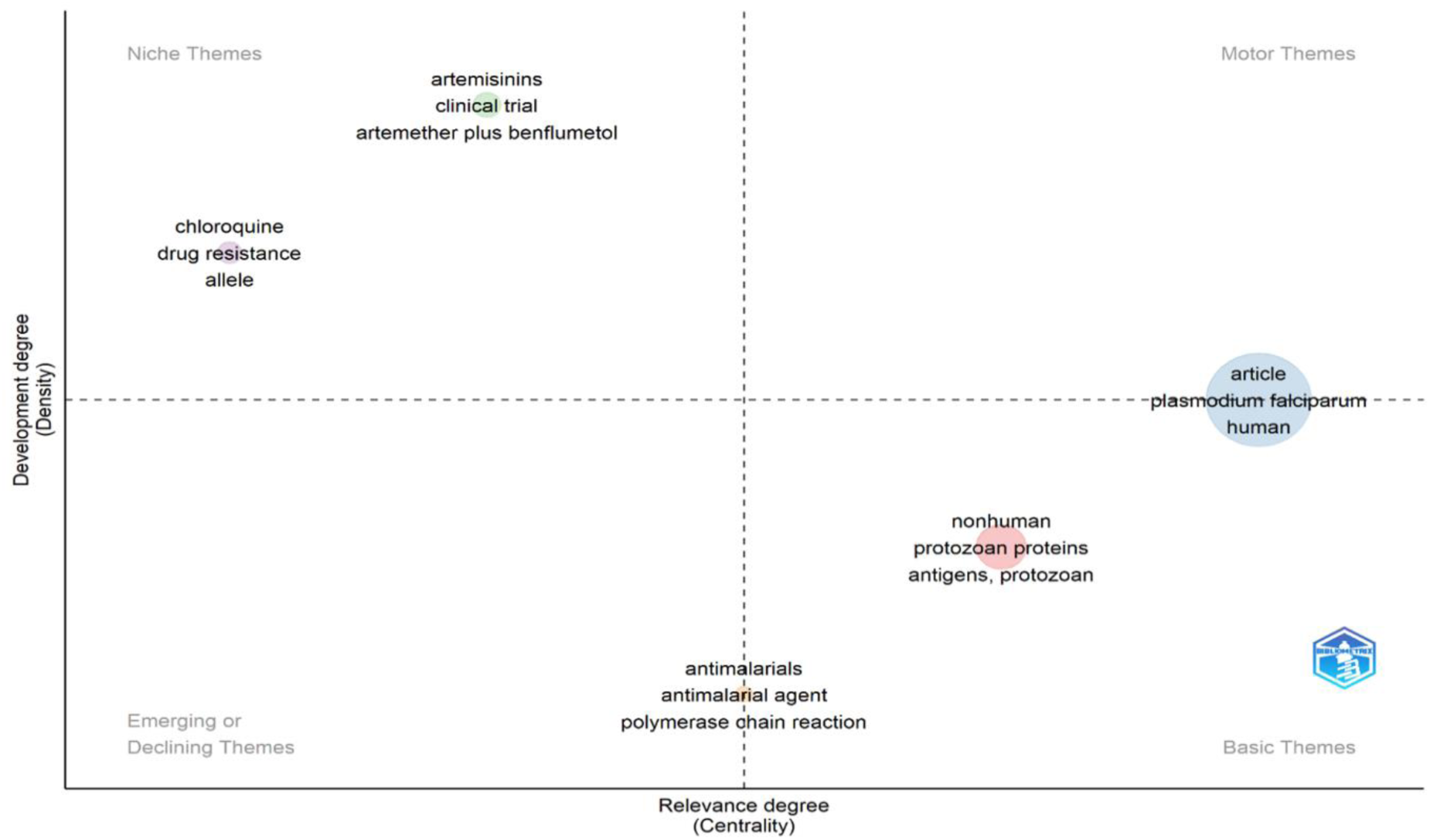
Thematic Map based on Authors Keyword on *Plasmodium falciparum* research in Africa.

### 3.6 Thematic Evolution of Authors’ Keywords used in Plasmodium falciparum Research in Africa

The thematic evolution of keywords in the research area of *Plasmodium falciparum* research in Africa was delineated into five distinct eras, consisting of 2000 to 2007, 2008 to 2012, 2013 to 2016, 2017 to 2020 and 2021 to 2024 as presented in Figure 14. During 2000 to 2007, 2008 to 2012, 2013 to 2016, 2017 to 2020, and 2021 to 2024, the major keywords in the order of occurrence were antimalarials and *Plasmodium falciparum* was prevalent in this era in research on *Plasmodium falciparum* in Africa.

**Figure 14:**
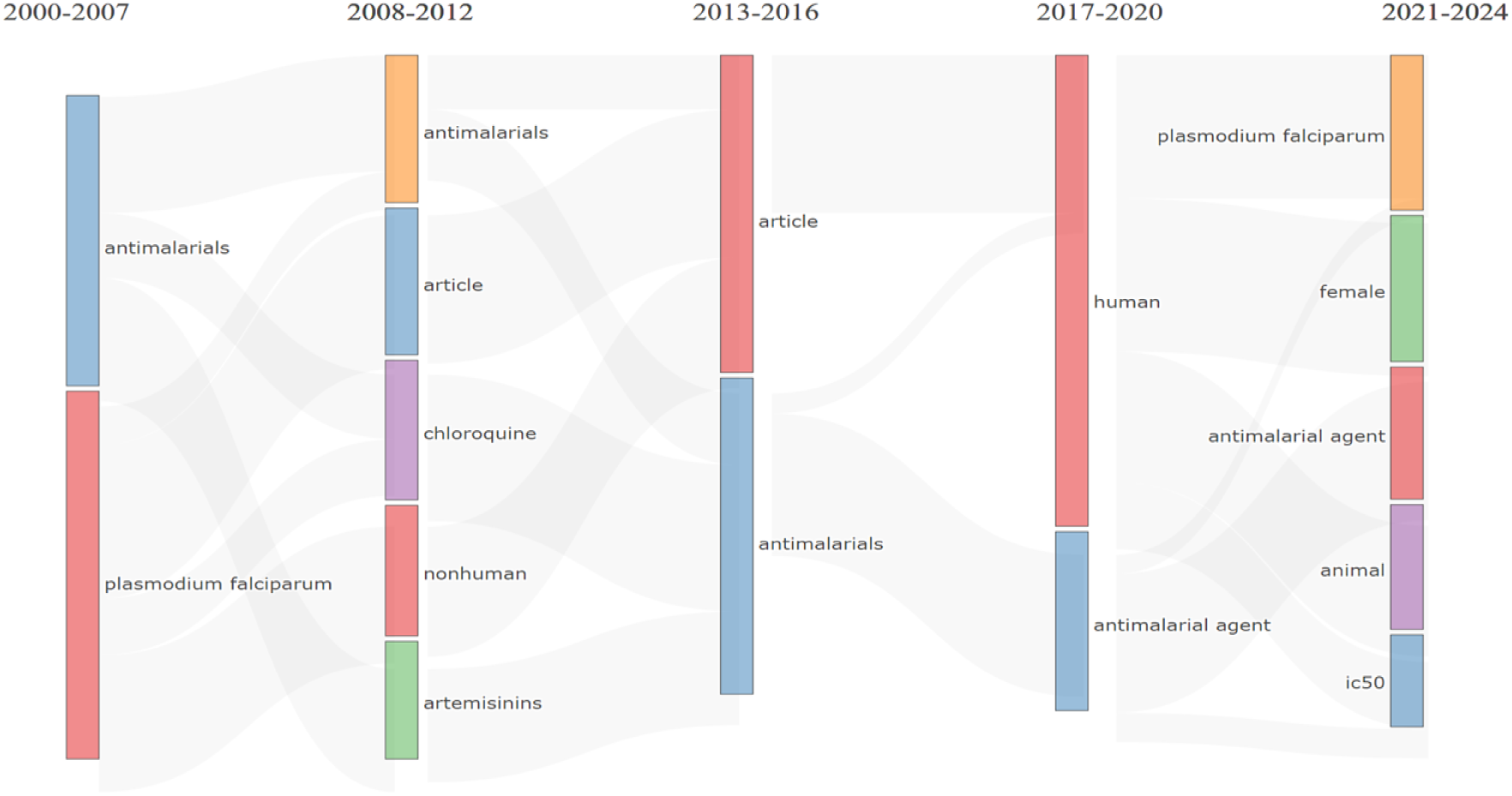
Thematic evolution of the Author’s Keywords used in *Plasmodium falciparum* research in Africa.

## 4.0 Co-occurrence network analysis

### 4.1 Co-occurrence-all keywords

Figure 15: presents the results on Keywords Co-occurrence Network in Global Research on *Plasmodium falciparum* research in Africa from 2000 to 2024 from Scopus database. Each node represents a keyword, and the size of the node designates the occurrence of the keyword. The largest nodes represent the most frequently used keywords in *Plasmodium falciparum* research. The link connecting two keywords indicates the number of times they co-occur. The color scale shows the average publication year of each keyword, where the color coding for each keyword is defined by the average publication year of all publications with the keyword. The keyword co-occurrence network map consists of nodes and lines. Each node represents a keyword while the size of the node designates the number of documents. The line connecting two nodes represents co-occurrence between the two keywords. The shorter the line, the stronger the co-occurrence and vice versa. We used the VOSviewer software to conduct a keyword co-occurrence analysis of all keywords, authors’ keywords and indexed keywords that have been used in *Plasmodium falciparum* research in Africa

**Figure 15:**
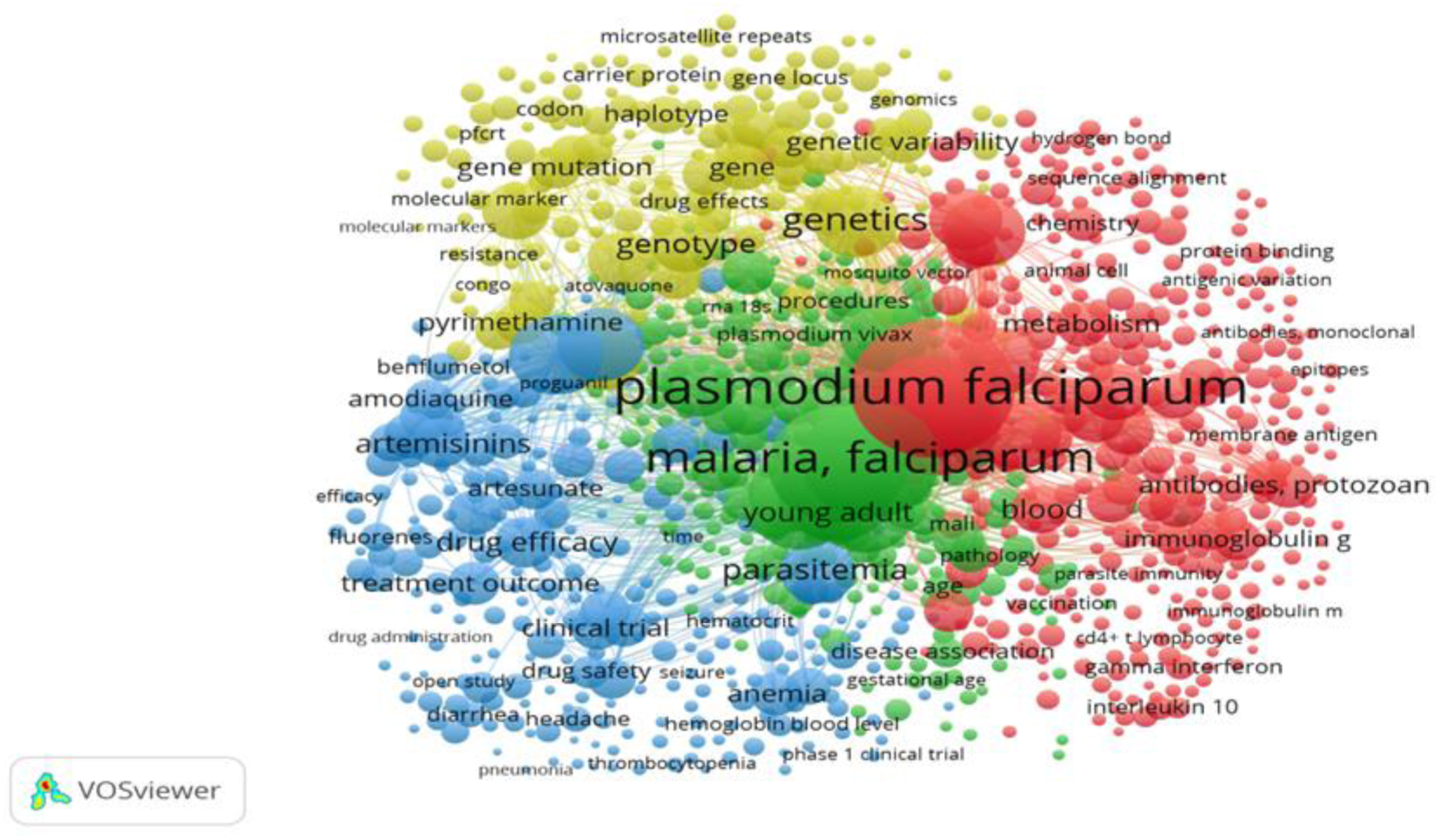
All Keywords Co-occurrence Network in Global Research on *Plasmodium falciparum* research in Africa.

**Figure 16:**
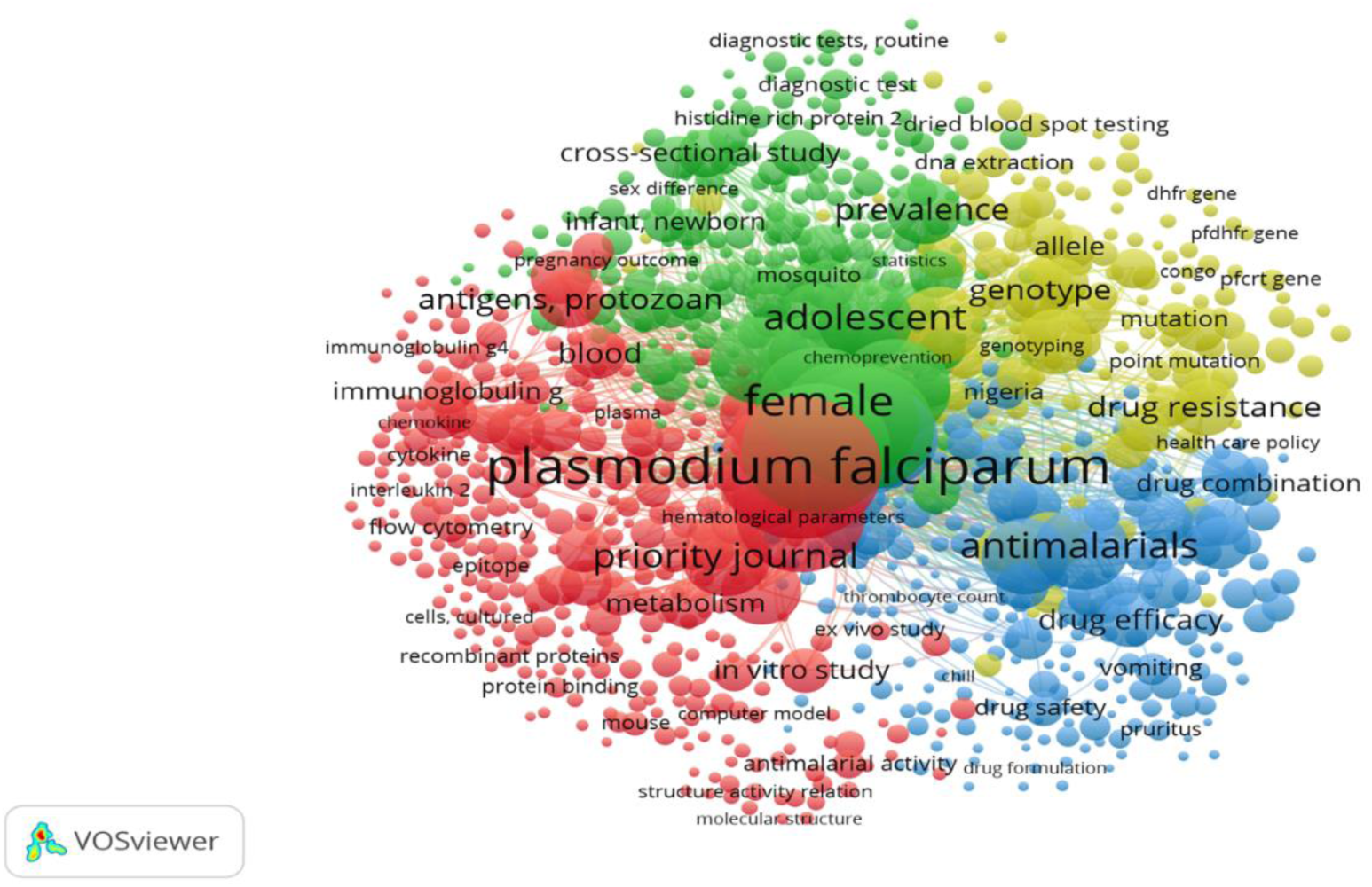
Index Keywords Co-occurrence Network in *Plasmodium falciparum* research in Africa.

**Figure 17:**
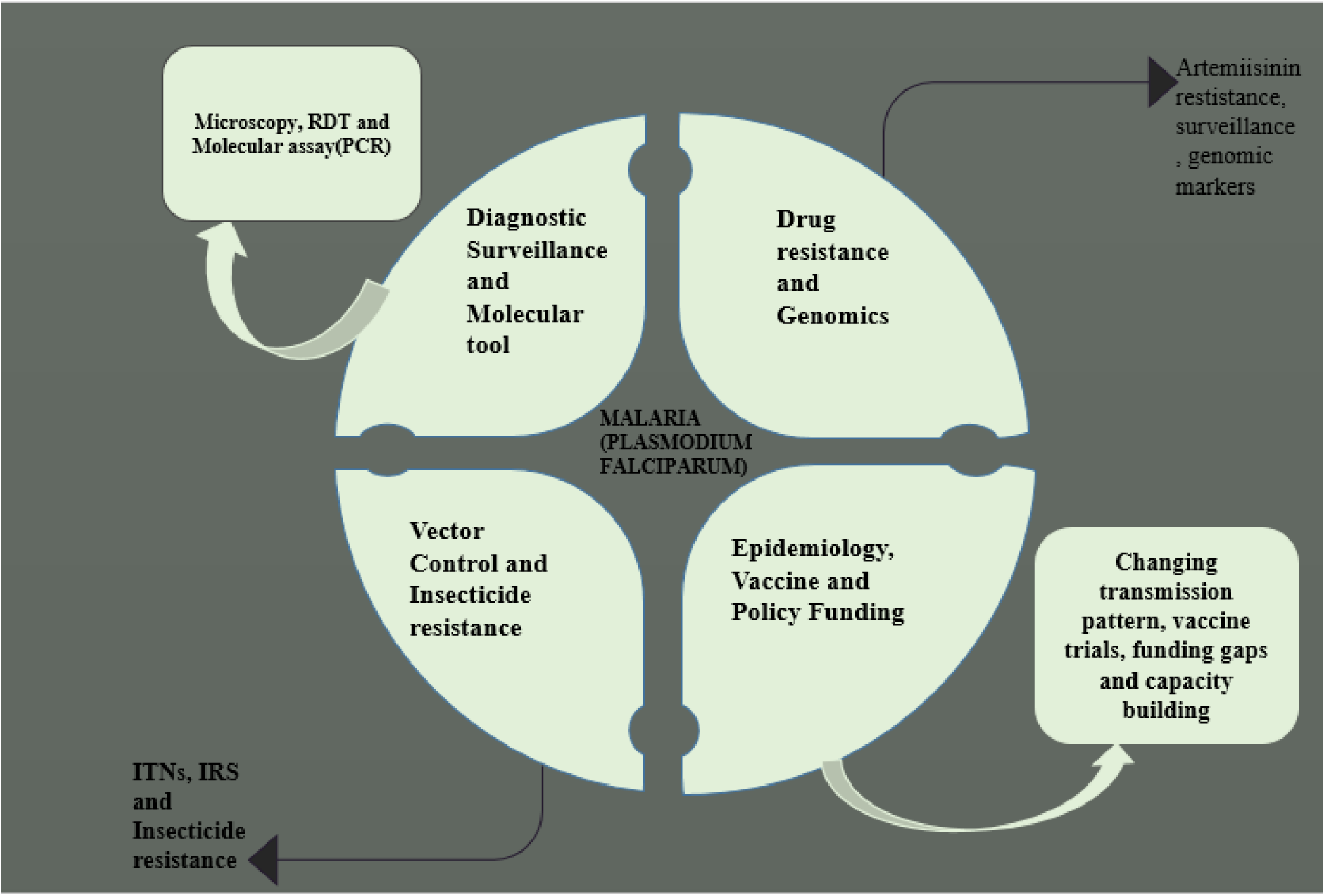
Schematic diagram and overview of *Plasmodium falciparum* research in Africa.

## 5.0: DISCUSSION

The bibliometric analysis focused on a review of studies to assess the type and amount of research on *Plasmodium falciparum* conducted in Africa from 2000 to 2024. The high volume of Primary Research (96%) indicates that *Plasmodium falciparum* research is an active and evolving field that suggests a focus on experimental studies, clinical trials, epidemiology research, drug resistance mechanisms, vaccine development, and new treatment strategies. Also, the amount of data generated led to discoveries in malaria pathogenesis and control. This distribution suggests a strong emphasis on primary research rather than literature synthesis. The 4% review might suggest that fewer efforts are being dedicated to consolidating existing research, and it could indicate a gap in comprehensive literature synthesis that analyses and contextualizes findings across different studies. A lack of systematic reviews and meta-analyses might slow down the application of research insights into clinical and public health strategies (Singh, 2017).

*Plasmodium falciparum* is the parasite responsible for the most severe form of malaria, evolves rapidly, developing resistance to both antimalarial drugs and insecticides, which poses a significant challenge to malaria control and elimination efforts. Drug resistance is primarily driven by mutations in key genes such as pfcrt, pfmdr1, dhfr, dhps, and kelch13, which confer resistance to widely used drugs like chloroquine, sulfadoxine-pyrimethamine, and artemisinin-based therapies; these mutations can arise independently in different regions or spread from a few resistant lineages across endemic areas, often in response to drug pressure and changes in treatment policies (Wicht *et al*., 2020;Thaloengsok *et al*., 2024) The evolution of resistance is further complicated due to accelerated resistance to multiple drugs (ARMD), where certain parasite populations acquire resistance to new drugs at high frequencies, raising concerns about the genetic mechanisms underlying this adaptability. Insecticide resistance in Anopheles mosquitoes, the primary malaria vectors, not only reduces the effectiveness of vector control tools but also increase the mosquitoes’ capacity to transmit *P. falciparum*, as resistance-associated physiological changes may enhance parasite infection and growth rates (Adams *et al*., 2022). Recent large-scale genomic studies and systematic in vitro evolution experiments have identified recurrently mutated genes and mutation patterns that predict resistance, offering new avenues for surveillance and drug development (Luth *et al*., 2024). Given the dynamic and region-specific nature of resistance evolution, continuous experimental research, systematic reviews, and meta-analyses are essential to synthesize findings, inform policy, and guide the development of standardized treatment and control strategies, ultimately reducing duplication of efforts and improving malaria interventions (Wicht *et al*., 2020).

The blue line on yearly mean citations of scientific publications on *Plasmodium falciparum* indicates a steady decline from 2000 to 2024 Mean TCperArt. This could reflect a decrease in the average citation impact per article over time. A possible explanation might be increased competition or a proliferation of research in the field leading to citation dilution. The MeanTCperYear shows fluctuation with a general increase up to around 2016, suggesting that yearly outputs were gaining more recognition. However, post-2016 there is a sharp decline, which could signal reduced attention to or relevance of *Plasmodium falciparum* research in recent years. In the gray lines, the peak observed in the mid-2010s may indicate key publications or breakthroughs, followed by a subsequent decline.

The declining trends might suggest a shift in research priorities or funding from *Plasmodium falciparum* to other areas like other malaria species, vaccine development, or broader public health concerns. The decline in average citations per article may indicate an increase in publication volume without corresponding citation growth, potentially reflecting overall citation impact per publication. The post-2016 decline in mean citations may point to a challenge in maintaining global interest in African-focused *Plasmodium* research. Understanding these trends is critical for strategizing funding, collaboration, and innovation in malaria research. The graph underscores the need for continued efforts to keep *Plasmodium falciparum* relevant, especially in the African context where malaria remains a pressing public health issue.

Partnerships from different countries characterize the global collaboration network in *Plasmodium falciparum* research in Africa, thus facilitated by shared research interest in *Plasmodium falciparum* research in Africa in African countries like Nigeria, Kenya, South Africa and Ghana are actively contributing to malaria research with significant documents counts with emphasis on the need to address the malaria burden in Africa, where *Plasmodium falciparum* is prevalent. The inter-country collaboration involves developed nations such as the USA, UK and France. This collaboration is vital for access to funding, advanced technologies, and knowledge exchange. These countries also play a key role in funding, technical support, and leading collaboration on *Plasmodium falciparum* research in Africa. An important aspect of this study revealed that in this bibliometric analysis there was a very high level of collaboration. Based on a review of publications, our survey revealed a high rate of collaboration between an African country’s institution and foreign/international institutions (Head *et al*., 2017). Countries such as Nigeria, Kenya, South Africa, and Ghana are actively involved in malaria research. They contribute significantly to the body of knowledge on *Plasmodium falciparum*, emphasizing the need to address the malaria burden in Africa. Developed countries play a pivotal role by providing funding, technical support, and leading collaboration efforts. These partnerships facilitate access to advanced technologies and knowledge exchange, which are essential for advancing malaria research.Projects like the Pan-African Malaria Genetic Epidemiology Network (PAMGEN) and the *P. falciparum* Community Project illustrate the collaborative efforts. PAMGEN aims to create a network of African scientists working with global researchers to use genetics and genomics for malaria elimination. It involves sampling from several African countries, including Cameroon, Ethiopia, The Gambia, Ghana, Madagascar, Mali, and Tanzania. The collaboration focuses on developing new tools for malaria control, understanding genetic variations in parasites, and nurturing African leadership in research. These efforts are critical for addressing the evolving challenges of malaria in Africa, such as drug resistance and vector adaptation. Strengthening research capacity and international partnerships can lead to innovative solutions for malaria control and eradication. This includes addressing emerging threats such as drug resistance and improving surveillance systems (Nankabirwa *et al*., 2022).

The steady decline from 2000 to 2024 suggests a decrease in the average citation impact per article over time. This could be due to increased competition or a proliferation of research in the field, leading to citation dilution. As more researchers enter the field, the average impact of individual articles may decrease because citations are spread across a larger number of publications. The fluctuation with a general increase up to around 2016 indicates that yearly outputs were gaining more recognition during that period. This could be attributed to significant advancements or breakthroughs in *Plasmodium falciparum* research, such as improved understanding of drug resistance or new control strategies. The sharp decline after 2016 might signal reduced attention to or relevance of *Plasmodium falciparum* research in recent years. This could be due to several factors, including shifts in global health priorities or the perception that malaria control efforts are becoming more effective, thus reducing the urgency for new research. The peak observed in the mid-2010s, as indicated by the gray lines, may correspond to key publications or breakthroughs. This period saw significant investments in malaria control and research, which could have led to notable findings and increased citations. The subsequent decline could reflect a normalization of interest following these major developments. Overall, these trends suggest that while *Plasmodium falciparum* research remains important, its citation impact and recognition may be influenced by broader trends in global health research and funding priorities. Despite these fluctuations, the ongoing challenge of malaria in Africa, particularly with drug resistance and transmission dynamics, underscores the need for continued research and innovation in this field (Amimo *et al*., 2020; D’Acremont *et al*., 2010; Noor *et al*., 2014).

Malaria being most prevalent in Africa, Kenya, Ghana, Niger, Nigeria and other African countries were African countries among the Top 20 countries with the highest publication outputs and coauthorships with Western countries. In its 2021 report, the WHO African Region indicates a disproportionately high share of the global malaria burden in Africa, accounting for more than three-quarters of the malaria burden (WHO, 2023). Similarly, children under the age of 5 accounted for about 80% of all malaria deaths in Africa, particularly in the SSA region (WHO, 2023). The Top 3 countries (United States, United Kingdom, and France), accounting for about three-quarters of the publication outputs, conducted most of the studies in collaboration with African countries. This corresponds to the number of malaria research publications, of which the United States and the United Kingdom dominated over half of the output from 1970 to 2022 (Aydemir *et al*., 2024). Nevertheless, the need for malaria research centers in Africa remains pertinent to promoting malaria research and successfully eradicating malaria globally.

The networks and collaborative efforts in studying *Plasmodium falciparum* highlight the importance of interdisciplinary approaches to tackle malaria comprehensively. These networks help identify dominant and emerging topics by analyzing keyword trends, which can guide African research institutions in aligning their focus areas with global priorities. This alignment ensures that their contributions remain impactful and relevant in addressing region-specific issues, such as malaria-endemic zones. This network studies the genetic diversity of *P. falciparum*, including molecular markers associated with resistance to artemisinin. It collaborates with institutions like the Wellcome Trust Sanger Institute to generate standardized data (Ndong Ngomo *et al*., 2023). Analyzing trends in research keywords helps identify emerging topics and areas of focus that are globally relevant. This ensures that African research institutions can align their research with international priorities, making their contributions more impactful (Apinjoh *et al*., 2019). The networks highlight opportunities for collaboration between African and international researchers. This collaboration is particularly important for addressing region-specific challenges, such as managing malaria in endemic zones (Apinjoh *et al*., 2019; Ndong Ngomo *et al*., 2023).

Basic research serves as the foundation for developing essential tools for malaria control, including drugs, insecticides, vaccines, and diagnostics. For instance, studies on phospholipid metabolism have led to the identification of novel antimalarial compounds (Ntoumi *et al*., 2004). Research on human and parasite gene polymorphisms provides valuable insights into the pathogenesis of severe malaria, aiding patient management and policy decisions (Ntoumi *et al*., 2004). Scientific investigations into antimalarial drug and insecticide resistance are crucial for sustaining effective control measures. Networks such as the Anti-malarial Drug Resistance Network (ADRN) support this research (Ntoumi *et al*., 2004). Additionally, African scientists are increasingly taking the lead in malaria research, ensuring that solutions are tailored to local contexts. Their leadership fosters trust and enhances the effectiveness of interventions (Thellier *et al*., 2024). Collaborative efforts between African and international researchers have further strengthened malaria research, facilitating large-scale studies and advancing technologies such as gene drive research (Thellier *et al*., 2024). Recent studies have shown that combining seasonal malaria vaccination with chemoprevention can significantly reduce malaria cases and deaths among African children, presenting a promising control strategy (Chandramohan *et al.,* 2021). Additionally, research has reinforced the effectiveness of long-lasting insecticidal nets (LLINs) and indoor residual spraying (IRS) in reducing malaria transmission. However, maintaining these interventions remains critical to prevent resurgence (Nankabirwa *et al*., 2022). Between 2000 and 2022, Africa experienced a 40% decrease in malaria incidence and a 60% reduction in mortality rates (Li *et al*., 2024). The Multilateral Initiative on Malaria (MIM), established in 1998, has played a significant role in increasing African involvement in malaria research, fostering collaboration and capacity building (Ntoumi *et al*., 2004). The availability of the malaria genome has also facilitated the discovery of novel drug targets and the development of new antimalarial compounds (Ntoumi *et al*., 2004). Initiatives such as the Program for Resistance, Immunology, Surveillance, and Modeling of Malaria (PRISM) in Uganda have contributed to a deeper understanding of malaria epidemiology and control interventions (Nankabirwa *et al*., 2022).

## 6.0: CONCLUSION AND RECOMMENDATION

This study highlights the significant progress in *P. falciparum* research in Africa, with increasing scientific output and strong international collaborations. However, persistent challenges such as drug and insecticide resistance, limited genomic surveillance, and socio-economic barriers to malaria control require further attention. Strengthening regional research collaborations, increasing funding for malaria research, and integrating multi-sectoral approaches will be critical in the fight against malaria. Future research should focus on next-generation vaccines, real-time surveillance systems, and community-based interventions to accelerate malaria elimination in Africa.

## Conflicts of Interest

The authors declare that they have no conflicts of interest.

## Source of funding

This review received no funding

## Notes

### Competing Interest Statement

The authors have declared no competing interest.

